# Epitope-Based Peptide prediction of vaccine against European bat lyssaviruses type 2 Glycoprotein G with Immunoinformatic Approach

**DOI:** 10.1101/2021.06.05.447201

**Authors:** Manal A. Gumaa, Abeer Babiker Idris, H. Bashir, Enas dk. Dawoud, Elhassn. Lina, Mohamed A. Hassan

## Abstract

**Objective:** European bat lyssaviruses (EBLV) type 2 is present in many European countries. Infection is usually seen in bats, the primary reservoirs of the viruses. Human deaths have been documented within few days following bat exposures. So, it is very useful to design an insilco peptide vaccine for European bat lyssaviruses type 2 virus using glycoprotein G as an immunogen to stimulate protective immune response

**Results:** B cell tests were conducted for Bepipred with 15 conserved epitopes, Emini surface accessibility prediction with 7 conserved epitopes in the surface and Kolaskar and Tongaonkar antigenicity tested with three conserved epitopes being antigenic. 357 conserved epitopes were predicted to interact with different MHC-1 alleles with (IC_50_) ≤500 while 282 conserved epitopes found to interact with MHC-II alleles with IC50≤ 1000. Among all tested epitopes for world population coverage the epitope **VFSYMELKV** binding to MHC11 alleles was 97.94% and it found to bind 10 different alleles that indicate strong potential to formulate peptide vaccine for lyssaviruses type 2 virus. To the best of our knowledge this is the first study to propose peptide vaccine for European bat lyssavirus type 2.

## Introduction

Rabies is one of most infectious diseases which is responsible for more than 50,000 human death every year, most of them are children[1, 2]. Lyssaviruses are single-stranded RNA viruses in the family Rhabdoviridae, genus Lyssavirus. Which has 12 species within the genus Lyssavirus, including the classical rabies virus and other closely related lyssaviruses such as the Australian bat lyssavirus (ABLV) and European bat lyssaviruses. Human infection can occur through scratching or bitting the skin, or through the contact with the mucosal surface. Human cases of rabies infection occur after animal bites or scratches, Aerosol transmission has never been documented in the natural environment but there were some cases of human infections due to animal licking of wounds or mucous membranes [3, 4].

First infection with bat rabies was reported in 1954. Then, more than 800 cases have been reported in different countries due to European Bat Lyssavirus Type-1 (EBLV-1) and Type-2 (EBLV-2)[3]. Some of bat species carry the viruses from more than 30 known species of bats which have been protected under the Agreement on the Conservation of Population of European bats[5, 6]. EBLVs are host-specific to specific bat species like Serotine bat (Eptesicus serotinus) which is mainly affected by EBLV1, while EBLV2 are most affected by mouse-eared bats (Myotis spp.) [7, 8].

The effect of current rabies vaccines against infection with other lyssaviruses has a critical rule in public health[9, 10]. The vaccines should give a significant cross-reactivity for African and Eurabian lyssaviruses with genetic modification[11]. Challenge studies in animal vaccination models suggest that there is protection provided by rabies vaccines against both the EBLVs, ABLV and some of the recently identified Asian lyssaviruses. LBV infection in rabies-vaccinated companion animals has highlighted the lack of protection against non-rabies lyssaviruses following vaccination. Further, pre- and postexposure vaccination failed to prevent disease and death in an animal model of (5.89%) of rabies infection [11, 12]. These factors suggest that more cross-reactive vaccine formulations may be necessary in areas where a threat to the human population comes from non-rabies lyssaviruses. Recent advances in the antigenic characterization of different lyssaviruses may also aid future cross-reactive vaccine design [11, 13, 14]. Genetic analysis of glycoprotein sequences from viruses circulating in both bat and terrestrial carnivore species suggests that host switching of lyssaviruses from bats to other mammals has occurred repeatedly and successfully, and Most viruses isolates from European bats was belonged to the RABV-serotype according to the antigenic typing studies using monoclonal antibodies[15, 16]. Then, the genotypes of EBLV-1 and EBLV-2 were identified according to their nucleotide and amino acid sequence and all were subdivided into two lineages[17, 18] [19]. Therefore, according to the genotypic differences, prevalence of infection and phenotypic differences may occur [19]. Studies with animal models, pre- and postexposure vaccination cannot avoid infections and death. So, for people who come from nonrabies lyssaviruses places they should take a reactive vaccine. Furthermore, to improve cross-reactive vaccine design. the studies should include an antigenic characterization of lyssaviruses[20, 21].

To our knowledge there is no peptide vaccine designed for European bat viruses, so it is very useful to design peptide vaccine with powerful immunogenic and minimal allergic effect. The aim of this study was to design an insilco peptide vaccine for European bat lyssaviruses type 2 virus using glycoprotein G as an immunogen to stimulate protective immune response.

## Main Text

### 2. Materials and methods

#### 2.1. Retrieval of envelope glycoprotein sequence

European bat lyssavirus 2 virus envelope glycoprotein sequences were obtained from NCBI (http://www.ncbi.nlm.nih.gov/protein/) database in December 2016. A total of 9 strains retrieved were collected from Europe; accession numbers of retrieved strains and their date and area of collection are listed in Table S1, Supplementary Material File.

#### 2.2. Conserved regions determination

Retrieved sequences were subject to multiple sequence alignment (MSA) using (Clustal W) as implemented in the BioEdit program, version 7.0.9.0[22] [23].

#### 2.3. Potential B cell epitopes prediction

The intent of the B cell epitope is to activate the B cell to produce the antibody specific for it (primary humoral response) or to convert the naive B cell into a memory B cell and make it ready to synthesize specific antibody in a second encounter [24].

The antigenic determinants of B-cell epitope are characterized by both being hydrophilic, accessible and in a flexible region of an immunogen [22, 24]. Thus the following implemented analyses were done computationally from the IEDB (http://www.iedb.org/) analysis resource [25].

➢ Bepipred Linear Epitope Prediction: It predicts the location of linear B-cell epitopes using a combinatorial algorithm made by combining the predictions of a hidden Markov model and the propensity scale by Parker *et a*l., thus it performs significantly better than any of the other methods [26]. Bepipred tool was used from the conserved region with a default threshold value of 0.031
➢ Emini surface accessibility scale : It was applied in the conserved region with a default threshold value1.00 [27].
➢ Kolaskar and Tongaonkar antigenicity scale: It was used to determine the antigenic sites with a default threshold value of 1.041 [28].

#### 2.4. T Cell Epitope Prediction

##### 2.4.1. CD8+ T-cell epitopes prediction

Several steps are required in the presentation of cytotoxic T cell epitopes. Although there are numerous algorithms have been developed to predict each of these steps in the CD8+ epitopes – MHC I binding pathway, a number of methods have even combined these steps to give an incorporated prediction of potential T cell epitopes [29, 30].

For the prediction of peptides binding to MHC-1, we used an artificial neural network (ANN) prediction method at Immune Epitope Database (IEDB) and predicted IC50 values for peptides binding to specific MHC molecules was less than 100 nm [31]. Prior to the prediction, peptide length was set to 9 amino acids and only frequently occurring alleles, that occur in at least 1% of the human population or allele frequency of 1% or higher, were selected, only those alleles [32]. Conserved epitopes having IC50 value less than 100 were considered as potential CD8+ T-cell epitopes and were selected for further analysis.

##### 2.4.2. CD4+ T-cell epitopes prediction

The ability of MHC class II groove to bind peptides with different lengths makes binding inaccuracy [33]. MHC-II binding predication was done by using NN-align method under MHC II binding prediction tool in IEDB analysis resource, which uses simultaneous identifies the MHC class II binding core epitopes and binding affinity [34]. Human allele references set were used. The conserved predicted CD4+ T-cell epitopes having higher binding affinity to interact with alleles at IC50 < 1000 were chosen for further analysis.

#### 2.5. Population coverage analysis

All MHC class I and MHC class II predicted T cell epitopes were assessed for population coverage against the whole world population by using the IEDB population coverage calculation tool [32].

#### 2.6. 3D structure modeling (homology modeling)

European bat lyssavirus 2 virus envelope glycoprotein 3D structure was obtained by RaptorX Property (http://raptorx2.uchicago.edu/StructurePropertyPred/predict/), which is a web server predicting structure property of a protein sequence without using any template information. This server is ranked 1st in the past 3 months in the fully-automated blind test CAMEO [35]. UCSF Chimera (version 1.8) was used to visualize the 3D structure, Chimera currently available within the Chimera package and available from the chimera web site (http://www.cgl.ucsf.edu/cimera) [36]. We obtained the 3D structure for further verification of the service accessibility and hydrophilicity of predicted B lymphocyte epitopes, as well as to identify all predicted T cell epitopes in the structural level.

**Table 1:** Accession numbers of retrieved strains and their date and area of collection

| GenBank Protein Accession No. | Country | Date of collection |
| --- | --- | --- |
| YP_001285396* | UK | 05-DEC-2006 |
| ABZ81190 | France | 19-NOV-2007 |
| ACA57832 | UK | 19-DEC-2007 |
| ABO65251 | UK | 05-DEC-2006 |
| AGG81481 | Finland | 05-JUN-2012 |
| AGG81486 | Finland | 05-JUN-2012 |
| AMR44682 | France | 24-FEB-2016 |
| ABZ81190 | France | 19-NOV-2007 |
| ACA57832 | UK | 19-DEC-2007 |

\* Reference strain

### 3. Result

#### 3.1. Potential B cell epitope prediction

The reference sequence of European bat lyssavirus 2 envelope glycoprotein was subjected to Bepipred linear epitope prediction, Emini surface accessibility and Kolaskar and Tongaonkar antigenicity prediction tools in IEDB, that predict the probability of specific regions in the protein to be potential B cell epitopes, being accessible, being immunogenic and being in a hydrophilic region.

Regarding Bepipred Linear Epitope Prediction 15 conserved epitopes were found to have the potential to evoke the B cell response. In Emini surface accessibility scale, 7 of them were found to satisfy the threshold value, 1.00. They were 83VVTEAETYTN92, 104KHFRPSP110, 112ACRDAYSWKTAGDPRYEESLHNPYPDSHWL141, 282FHSD285, 342MEADAHYK349, 369RCYPH373 and 440PDVQKQISG448. While in Kolaskar and Tongaonkar antigenicity scale, 3 of them gave score above threshold 1.041. They were 49VEDEGCT55, 111ACRDAYSWKTAGDPRYEESLHNPYPDSHWL141 and 440 PDVQKQISG448. However, 2 conserved B cell epitopes from all retrieved strains were found to satisfy the all three scales of B cell epitope prediction tools. The linear epitope 112ACRDAYSWKTAGDPRYEESLHNPYPDSHWL141 showed the highest score in accessibility and antigenicity test followed by 440 PDVQKQISG448. The result is summarized and illustrated in Figure (S1, S2, S3) and Table S2, Supplementary Material File. Proposed epitopes positions in the structural level are shown in (Figure S4-S5), Supplementary Material File.

#### 3.2. Potential T-cell epitope prediction

##### 3.2.1. CD8+ T-cell epitope prediction

The reference sequence of European bat lyssavirus 2 virus envelope glycoprotein was analyzed using IEDB MHC-1 binding prediction tool to predict T cell epitopes suggested interacting with most frequently occurring alleles of MHC Class I alleles. ANN based MHC 1 binding prediction tool with IC50 < 100; 357 conserved peptides were predicted to interact with different MHC-1 alleles. The epitope 254LMDGSWVSI262 and 440PDVQKQISG448 had higher affinity to interact with 4 alleles followed by FLNGKCSGV, RLMDGSWVS, VLIPEMQSA and VLSALLSAL which interact with 3 alleles. Epitopes and their corresponding alleles are illustrated in (Table S3), and proposed epitopes are shown in the structural level in Figure (S6-S7), Supplementary Material file.

##### 3.2.2. CD4+ T-cell epitope prediction

By using NN-align method under MHC II binding prediction tool in IEDB analysis resource with IC50 < 1000; 282 conserved peptides were predicted to interact with different MHC-II alleles. The peptide (core) 59VFSYMELKV67 and 57LTVFSYMEL65 had high affinity to interact with 10 alleles (HLA-DPA1*01, HLA-DPA1*01:03, HLA-DPA1*02:01, HLA-DPA1*03:01, HLA-DPB1*01:01, HLA-DPB1*02:01, HLA-DPB1*04:01, HLA-DPB1*04:02, HLA-DPB1*05:01, HLA-DRB1*15:01) and (HLA-DPA1*01, HLA-DPA1*01:03, HLA-DPA1*02:01, HLA-DPA1*03:01, HLA-DPB1*01:01, HLA-DPB1*02:01, HLA-DPB1*04:01, HLA-DPB1*04:02, HLA-DQA1*01:02, HLA-DQB1*06:02), respectively. The result is listed in (Table: S4) and proposed epitopes are shown in the structural level in Figure (S8-S9), Supplementary Material file.

#### 3.3. Population coverage analysis

All MHC class I and MHC class II predicted T cell epitopes (especially whose with high affinity binding epitopes and can bind to different set of alleles) were selected for population coverage analysis. The result is summarized in Table S5, Supplementary Material file.

### 4. Discussion

Prevention of human rabies is accomplished by controlling rabies in domestic and wild animals, including the use of vaccination programs. The usefulness of human rabies vaccines is hampered by high cost, complicated vaccination regimens and lack of compliance, especially in areas of Africa and Asia where human rabies infections are endemic [37]. According to any studies approved that the prophylactic and vaccinations still the important way to avoid human rabies infection[38]. The vaccine should be improved to avoid the complications of multi doses and it is highly costs may affect the people to get the vaccine in many countries [39]. Study done in UK showed that the vaccines cannot inhance the CD4+ T-cell to activate the immune response against the virus when they reduce the injections number with the same inoculum (<0.5 IU/ml) in animal model [40]. Another Study detect the effectiveness of Post exposure rabies vaccine (SRV9) using one dose of vaccine by an animal inoculation, show that the vaccine is effective in about 70 % of rates which was injected in the first day of infection [41].

In this study we selected different peptides that could be recognized by B cell and T cell to produce antibodies against European bat lyssavirus type 2. This peptide vaccine was expected to be more antigenic and less allergic than the conventional biochemical vaccines that based on live attenuated virus. Pre- and postexposure vaccination failed to prevent disease and death in an animal model of European bat lyssavirus infection. These factors suggest that more cross-reactive vaccine formulations may be necessary in areas where a threat to the human population comes from non-rabies lyssaviruses. Recent advances in the antigenic characterization of different lyssaviruses may also aid future cross-reactive vaccine design

The reference sequence of European bat lyssavirus type 2 glycoprotein (GP) was subjected to Bepipred linear epitope prediction test, Emini surface accessibility test and Kolaskar and Tongaonkar antigenicity test in IEDB, to determine the binding to B cell, being in the surface and to test the immunogenicity respectively. For Bepipred test there was 15 conserved epitopes that have the binding affinity to B cell (table 2). While there are 7 epitopes predicted on the surface according to Emini surface accessibility test. Three epitopes being antigenic as detected by Kolaskar and Tongaonkar antigenicity scale, while only two conserved B cell epitopes (112ACRDAYSWKTAGDPRYEESLHNPYPDSHWL141and 440 PDVQKQISG448) found to overlap all three performed B cell epitope prediction tool tests.

Also the reference sequence of European bat lyssavirus type 2 envelope glycoprotein G was analyzed using IEDB MHC-1 binding prediction tool to predict T cell epitope suggested to interact with different types of MHC I alleles. Based on Artificial neural network (ANN) with half-maximal inhibitory concentration (IC_50_) ≤500; 23 conserved peptides were predicted to interact with different MHC-1 alleles. The peptide 254LMDGSWVSI262 had higher affinity to interact with 4 alleles (HLA-A*01:01, HLA-A*02:01, HLA-A*02:06, HLA-A*32:01), and the peptide 410SVIPLMHPL418 that binds to the same numbers of alleles (HLA-A*01:01, HLA-A*02:01, HLA-A*02:06, HLA-A*68:02).These two peptides could be thought about as a possible peptide vaccine for European bat lyssavirus type 2.

The reference glycoprotein (GP) strain was analyzed using IEDB MHC-II binding prediction tool based on NN-align with half-maximal inhibitory concentration (IC_50_) ≤1000; there were 282 conserved predicted epitopes found to interact with MHC-II alleles.The peptide (core) 59VFSYMELKV67 and 57LTVFSYMEL65 had high affinity to interact with the highest number of MHC11 alleles (10 alleles).

World population coverage results for total epitopes binding to MHC1 alleles was 58.53% and the most promising peptides was LMDGSWVSI and SVIPLMHPL with world population coverage 56.89% and 55.47% respectively with total HLA hits 4. The world population coverage results for all epitopes that have binding affinity to MHC11 alleles was 98.65% while world population coverage of the most promising two epitopes VFSYMELKV and LTVFSYMEL was 97.94% and 97.52% respectively with HLA hits 10.

### 7. Limitations

## 7. Declarations

### Ethics approval and consent to participate

The study protocol was approved by the National Health Research Ethics Committee authorization.

### Consent for publication

Not Applicable

### Availability of data and material

The datasets used and analysed during the current study available on NCBI (http://www.ncbi.nlm.nih.gov/protein/) database.

### Author’s contributions

The authors confirm contribution to the paper as follows: analysis and interpretation of results: M. Gumaa; verified the analytical methods: A. Babiker; study conception and design: N. Bilal; supervised the findings of this work: M. Hassan. All authors reviewed the results and approved the final version of the manuscript.

## References

1. Jackson, A.C.J.J.o.n., Rabies virus infection: an update. 2003. 9(2): p. 253–258.

2. Nel, L.H.J.E.i.d., Discrepancies in data reporting for rabies, Africa. 2013. 19(4): p. 529.

3. Fooks, A.R., et al., Case report: isolation of a European bat lyssavirus type 2a from a fatal human case of rabies encephalitis. 2003. 71(2): p. 281–289.

4. Calisher, C.H., J.A.J.T.m. Ellison, and i. disease, The other rabies viruses: The emergence and importance of lyssaviruses from bats and other vertebrates. 2012. 10(2): p. 69–79.

5. Mitchell-Jones, A.J., et al., The atlas of European mammals. Vol. 3. 1999: Academic Press London.

6. Vos, A., et al., European bat lyssaviruses—an ecological enigma. 2007. 9(1): p. 283–296.

7. Stantic-Pavlinic, M.J.E., Public health concerns in bat rabies across Europe. 2005. 10(11): p. 5–6.

8. Faber, M., et al., Immunogenicity and safety of recombinant rabies viruses used for oral vaccination of stray dogs and wildlife. 2009. 56(6-7): p. 262–269.

9. Müller, T., et al., Analysis of vaccine-virus-associated rabies cases in red foxes (Vulpes vulpes) after oral rabies vaccination campaigns in Germany and Austria. 2009. 154(7): p. 1081–1091.

10. Prager, K., et al., Vaccination strategies to conserve the endangered African wild dog (Lycaon pictus). 2011. 144(7): p. 1940–1948.

11. Velasco-Villa, A., et al., Successful strategies implemented towards the elimination of canine rabies in the Western Hemisphere. 2017. 143: p. 1–12.

12. Rota Nodari, E., et al., Rabies vaccination: higher failure rates in imported dogs than in those vaccinated in Italy. 2017. 64(2): p. 146–155.

13. Huang, F., et al., Efficiency of live attenuated and inactivated rabies viruses in prophylactic and post exposure vaccination against the street virus strain. 2015. 59(2): p. 117–24.

14. Li, J., et al., Postexposure treatment with the live-attenuated rabies virus (RV) vaccine TriGAS triggers the clearance of wild-type RV from the Central Nervous System (CNS) through the rapid induction of genes relevant to adaptive immunity in CNS tissues. 2012. 86(6): p. 3200–3210.

15. Schneider, L., J.J.C.t.i.m. Cox, and immunology, Bat lyssaviruses in Europe. 1994. 187: p. 207–218.

16. Bourhy, H., et al., Antigenic and molecular characterization of bat rabies virus in Europe. 1992. 30(9): p. 2419–2426.

17. Amengual, B., et al., Evolution of European bat lyssaviruses. 1997. 78(9): p. 2319–2328.

18. Davis, P.L., et al., The evolutionary history and dynamics of bat rabies virus. 2006. 6(6): p. 464–473.

19. Wu, X., et al., Development of combined vaccines for rabies and immunocontraception. 2009. 27(51): p. 7202–7209.

20. Horton, D.L., et al., Quantifying antigenic relationships among the lyssaviruses. 2010. 84(22): p. 11841–11848.

21. Banyard, A.C., et al., Bats and lyssaviruses, in Advances in virus research. 2011, Elsevier. p. 239–289.

22. Florian, D.D., et al., In silico modeling and immunoinformatics probing disclose the epitope based peptidevaccine against Zika virus envelope glycoprotein. 2014. 2(04): p. 44–57.

23. Hall, T.A. BioEdit: a user-friendly biological sequence alignment editor and analysis program for Windows 95/98/NT. in Nucleic acids symposium series. 1999. [London]: Information Retrieval Ltd., c1979-c2000.

24. Welling, G.W., et al., Prediction of sequential antigenic regions in proteins. 1985. 188(2): p. 215–218.

25. Vita, R., et al., The immune epitope database (IEDB) 3.0. 2015. 43(D1): p. D405–D412.

26. Larsen, J.E.P., O. Lund, and M.J.I.r. Nielsen, Improved method for predicting linear B-cell epitopes. 2006. 2(1): p. 1–7.

27. Emini, E.A., et al., Induction of hepatitis A virus-neutralizing antibody by a virus-specific synthetic peptide. 1985. 55(3): p. 836–839.

28. Kolaskar, A.S. and P.C.J.F.l. Tongaonkar, A semi-empirical method for prediction of antigenic determinants on protein antigens. 1990. 276(1-2): p. 172–174.

29. Lundegaard, C., M. Nielsen, and O.J.T.i.B. Lund, The validity of predicted T-cell epitopes. 2006. 24(12): p. 537–538.

30. Badawi, M.M., et al., Highly conserved epitopes of Zika envelope glycoprotein may act as a novel peptide vaccine with high coverage: immunoinformatics approach. 2016. 4(3): p. 46–60.

31. Buus, S., et al., Sensitive quantitative predictions of peptide-MHC binding by a ‘Query by Committee’artificial neural network approach. 2003. 62(5): p. 378–384.

32. Bui, H.-H., et al., Predicting population coverage of T-cell epitope-based diagnostics and vaccines. 2006. 7(1): p. 1–5.

33. Badawi, M.M., et al., Immunoinformatics predication and in silico modeling of epitope-based peptide vaccine against virulent Newcastle disease viruses. 2016. 4(3): p. 61–71.

34. Nielsen, M. and O.J.B.b. Lund, NN-align. An artificial neural network-based alignment algorithm for MHC class II peptide binding prediction. 2009. 10(1): p. 296.

35. Wang, S., et al., RaptorX-Property: a web server for protein structure property prediction. 2016. 44(W1): p. W430–W435.

36. Ferrin, U., Chimera a visualization system for exploratory research and analysis. 2004, J.

37. Rashid, A., K. Rasheed, and M.J.J.A.P.S. Akhtar, Factors influencing vaccine efficacy: a general review. 2009. 19: p. 22–25.

38. Franka, R., et al., Rabies virus pathogenesis in relationship to intervention with inactivated and attenuated rabies vaccines. 2009. 27(51): p. 7149–7155.

39. Blaney, J.E., et al., Inactivated or live-attenuated bivalent vaccines that confer protection against rabies and Ebola viruses. 2011. 85(20): p. 10605–10616.

40. Strady, C., et al., Immunogenicity and booster efficacy of pre-exposure rabies vaccination. 2009. 103(11): p. 1159–1164.

41. McGettigan, J.P.J.E.r.o.v., Experimental rabies vaccines for humans. 2010. 9(10): p. 1177–1186.

